# The impact of heterogeneous spatial autocorrelation on comparisons of brain maps

**DOI:** 10.1101/2024.06.14.598987

**Authors:** R Leech, JS Smallwood, R Moran, EJH Jones, N Vowles, D Leech, EM Viegas, FE Turkheimer, F Alberti, D Margulies, E Jefferies, A Alexander-Bloch, Y Zhou, B Bernhardt, F Váša

## Abstract

It is increasingly common to statistically compare brain maps to assess how spatially similar they are. However, statistical inference can be challenging due to the presence of spatial autocorrelation. Therefore, random permutation approaches based on null models are widely used to address this concern. Here, we show how that the presence of heterogeneity in the spatial autocorrelation across brain maps impacts the validity of statistical inference in common approaches for spatially correlated maps. Furthermore, we illustrate how a Bayesian spatial regression approach can be applied to compare functional and structural cortical brain maps, yielding valid statistical inferences even in the presence of heterogeneity. Explicitly modelling spatial properties provides more valid inferences about whole brain spatial maps allowing a wider and more sophisticated range of neurobiological questions to be answered about the relationship between brain maps than are possible with current approaches.

## Introduction

Over recent decades, there has been a growing realisation of the importance of the topographic organisation of the brain. One key method for investigating neural topography is to quantitatively compare spatial maps that capture different biological features. For example, recent studies have compared whole-brain spatial maps measured with fMRI at rest with a range of morphological, molecular and genetic whole-brain spatial patterns (Paquola et al., 2019; Váša et al., 2020; Hansen et al., 2022; Larivière et al., 2023; Wagstyl et al., 2024).

Statistical comparisons between brain maps are challenging, however, because of the presence of spatial autocorrelation. This phenomenon refers to spatially proximal measurements being more similar, in general (Burt et al., 2020; Shinn et al., 2022; Leech et al., 2024, 2023). In the presence of spatial autocorrelation, the significance of similarity between pairs of brain maps is likely to be inflated. In fact, p-values resulting from measures of statistical association such as Pearson’s or Spearman’s correlation coefficient assume that underlying datapoints are independent (and the number of datapoints directly translates into degrees of freedom for the correlation). This assumption is violated by spatial autocorrelation (Worsley et al., 1999; Alexander-Bloch et al., 2018; Váša and Mišić, 2022).

Approaches for addressing the consequences of spatial autocorrelation on comparisons of brain maps have predominantly involved the use of null models which aim to preserve spatial autocorrelation between the true and synthesised data (Alexander-Bloch et al., 2018; Burt et al., 2020; Markello and Mišić, 2021; Vos de Wael et al., 2020). One approach involves iterative randomisation and smoothing of brain maps to generate surrogate maps with randomized topography but preserved spatial autocorrelation (Burt et al., 2020, 2018). Another popular approach applies a spatial randomization or “spin” to a spherical projection of the data (Alexander-Bloch et al., 2018; Váša et al., 2018). Alternative approaches propose to permute spectral decompositions of the geodesic distance matrix (Vos de Wael et al., 2020), or to randomly rotate a spectral decomposition of cortical and subcortical surfaces into geometric eigenmodes (Koussis et al. 2024). These spatial null models can be used to benchmark the magnitude of the similarity between two brain maps against a null distribution corresponding to the similarity between one of the maps, and numerous surrogate instances of the other map (Váša and Mišić, 2022).

At the core of randomization approaches used in studies of brain maps is the assumption that autocorrelation across maps is spatially stationary, or homogeneous (see left panel in Figure 1). In other words, the data is assumed to come from a random distribution with statistical properties that do not change over space. However, evidence suggests that there is considerable spatial heterogeneity in autocorrelation across the cortex for functional MRI (Hayasaka et al., 2004; Leech and Leech, 2011; Li et al., 2015; Leech et al. 2023), and such findings are likely similar for other modalities (Paquola et al., 2023). If the spatial autocorrelation varies systematically across the brain – e.g. functional properties change slower over space in regions of primary cortex than in association cortex, see Leech et al., (2023) – then this may make it hard to draw valid statistical inferences regarding the similarity of brain maps.

**Figure 1.**
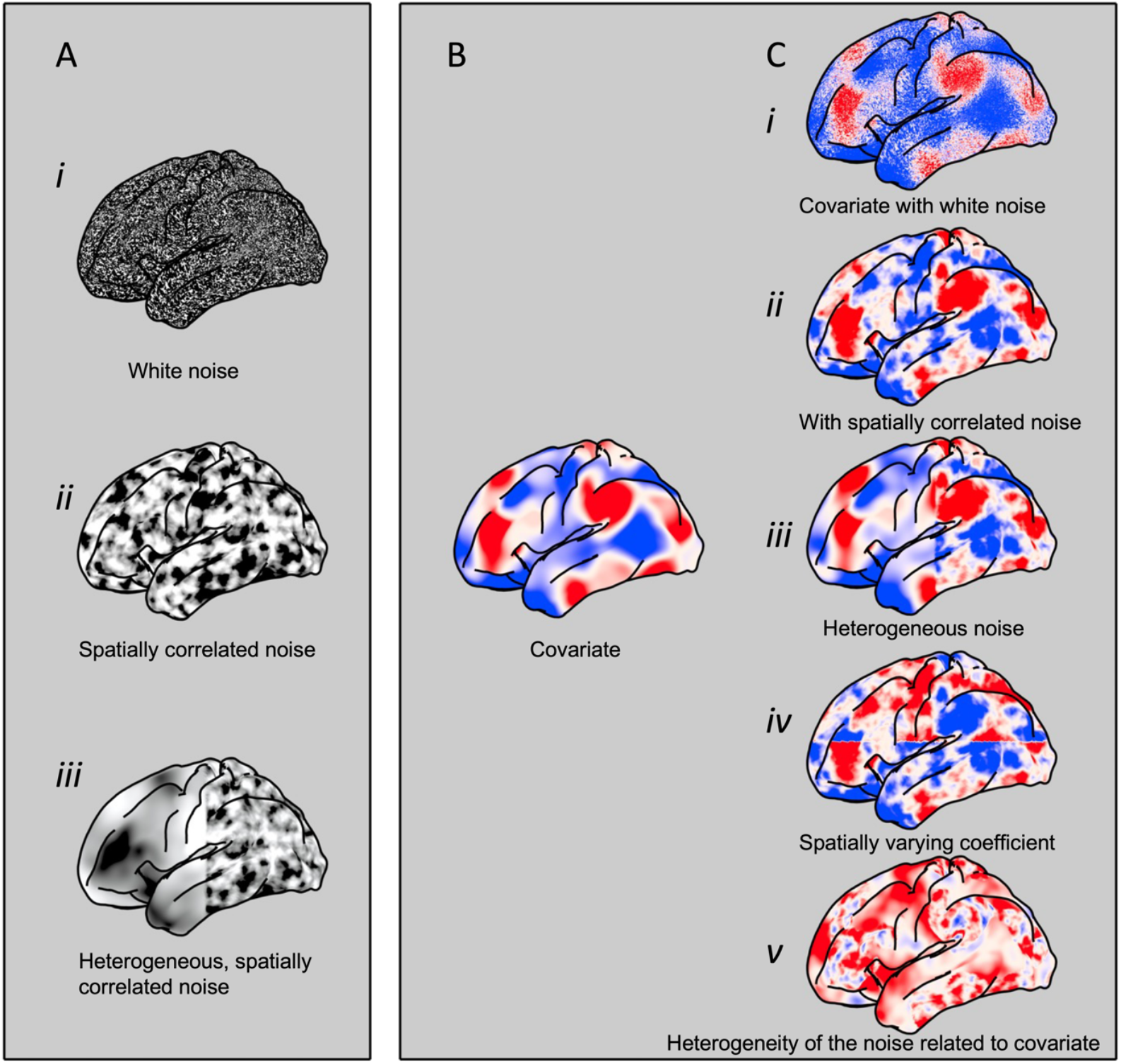
Heterogeneous spatial structure in brain maps. Left panel: A) Illustration of different types of spatial patterns across the cortex: **(i)** independently and identically distributed white noise, (ii) spatially correlated noise generated with a spatially homogeneous covariance structure; (iii) spatially heterogeneous noise, with a greater spatial noise scale anteriorly than posteriorly. Right panel: some of the potential relationships between two spatial maps: B) is a randomly generated map with spatially-correlated structure; C) are example illustrations of other maps which are related to the first map in different ways: (i) white noise and a consistent spatial relationship between the two maps; (ii) homogeneous spatially correlated noise and a homogeneous spatial relationship between the two maps; (iii) spatially heterogeneous spatial noise and a consistent spatial relationship between maps; (iv) homogeneous spatial noise and a spatially varying coefficient between the two maps (superior half has a positive relationship, inferior half has a negative relationship). (v) no spatially varying correlation, but spatially varying autocorrelated noise between the two maps.

There are many possible spatial processes that could be responsible for heterogeneity in spatial autocorrelation. For example, it could result from changes across space in scanner noise, motion artefacts or other measurement issues, or equally from neurobiological processes that have not been accounted for such as structural properties or topographical neurobiological organisation; e.g., in certain regions of cortex, signal from multiple sources are mapped in close proximity (Goldman-Rakic and Schwartz, 1982).

Spatial autocorrelation and heterogeneity across space can take multiple forms (**Figure 1**). For the case of a single brain map (Figure 1A) we see e.g., (i) a pattern with no spatial structure; (ii) homogeneous spatial autocorrelation across the brain; or (iii) a pattern with a spatially varying spatial autocorrelation structure (e.g., changing length scale of the pattern from the front to the back of the brain). The situation becomes more complicated when comparing one spatial map (e.g., **Figure 1B**) with another (illustrated in **Figure 1C**). Here, we can potentially have heterogeneity both in the measured pattern (within each brain map) and in the relationship between the two maps. To illustrate, the simplest case is Gaussian noise (*i*.*e*., no spatial autocorrelation) and a homogeneous correlation between the two maps across space (**Figure 1C i**); in this situation simple correlation or ordinary least squares regression is appropriate for comparing the two maps. The second situation (**Figure 1C ii**) is one where the spatial autocorrelation is homogeneous and there is a consistent relationship between the two maps across the brain; here, existing null permutation approaches are appropriate. The third illustration (**Figure 1C iii**) involves a heterogeneous spatial pattern and a consistent spatial correlation across the two maps. In this scenario some areas of cortex are consistently homogeneous while other are heterogeneous. The fourth example (**Figure 1C iv)** has homogeneous spatial autocorrelation of the noise, but the correlation between the maps varies over space (with a positive correlation coefficient in the posterior parts of the cortex and a negative correlation coefficient in the inferior parts). The final illustration (**Figure 1C v**) involves no spatial correlation between the two maps; however, the spatial autocorrelation in the second map varies depending on values of the first map. The final three hypothetical examples pose challenges for existing null model-based approaches for comparing spatial maps because simply permuting or randomizing the data may fail to capture systematic features of the true data, potentially leading to inappropriate statistical inference.

In this paper, we first illustrate that the assumption that spatial variation is homogeneous can lead to problems when statistically comparing spatial maps as is commonly performed in the neuroimaging field. To do so, we first quantify the extent of heterogeneity in spatial autocorrelation exhibited by multiple brain maps generated using different imaging modalities. Next, we use synthetic data to illustrate why such heterogeneity can lead to problems with statistical inference when comparing spatial maps using existing null model approaches.

The issues of heterogeneity when performing spatial correlation have been recognized in other fields, notably ecology, climatology and epidemiology (Daly et al., 1994). Recent theoretical and computational approaches have allowed for sophisticated modelling of spatial structure, including different forms of heterogeneity (as illustrated in **Figure 1**) even for large datasets. We use one such approach, based on stochastic partial differential equations, to explicitly model and compare potential spatial processes underlying brain maps (Lindgren and Rue, 2015). Using this approach, we show how statistical assessments between brain maps can be made while modelling different forms of spatial heterogeneity, using both synthetic and real neuroimaging data, and therefore, providing a framework to make more valid statistical inferences regarding macroscale properties of the cortex. We also highlight how alternative analysis approaches allow us to ask more sophisticated questions about spatial relationships between brain maps. In particular, the proposed approach enables estimation of a continuous spatially-varying relationship between maps (such as illustrated Figure1C iv), compared to prior approaches which only enable quantifying a single homogeneous measure of similarity.

## Results

### Heterogeneity in spatial autocorrelation across the cortex

We first consider several canonical maps of cortical organization from different imaging modalities obtained from *neuromaps* (Markello et al., 2022). For this example, we consider the T1w/T2w ratio (a proxy for intracortical myelin) and cortical thickness (Glasser et al., 2013), the principal functional connectivity gradient derived from resting-state fMRI (Margulies et al., 2016), and a map of evolutionary expansion from non-human primates and humans (Hill et al., 2010) (**Figure 2A**). For each map, we calculated the local Moran’s I statistic (Anselin, 1995; Moran, 1950) for each vertex using a geodesic distance matrix along the cortical surface. Moran’s I is calculated based on the pairwise distances and dissimilarities between all data points. It results in a statistic with values between −1 and 1 which indexes the magnitude and sign of spatial autocorrelation, while 0 indicates the absence of autocorrelation. Local Moran’s I calculates a similar metric but separately for each data point compared to all other points (See **Figure 2B** for an illustration). A high score on this metric reflects a high degree of spatial similarity in a local region of cortex.

**Figure 2.**
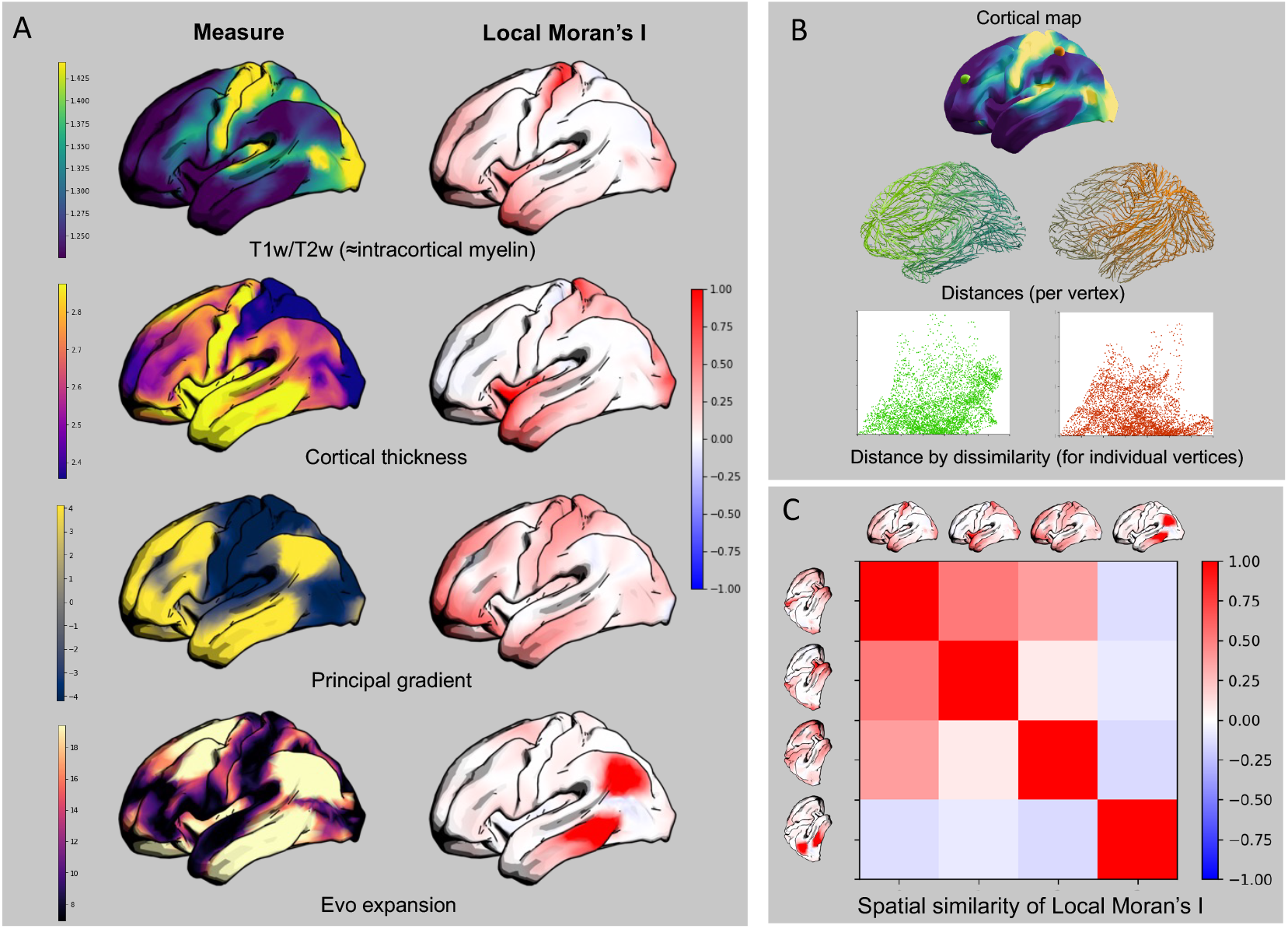
Regionally heterogeneous spatial autocorrelation. A) left: Four exemplar cortical maps: T1w/T2w ratio, cortical thickness, the first functional connectivity gradient, cortical expansion between macaque monkey and human. Right, the distribution of local Moran’s I across the cortex (measuring regional variability in spatial autocorrelation). B) Illustration of calculating local Moran’s I for two illustrative vertices from the T1w/T2w cortical map, combining dissimilarity to all other vertices with the vector of distances to other vertices. C) The spatial similarity in the distribution of local Moran’s I across maps, quantified using Spearman’s ρ.

In **Figure 2A**, it can be seen that positive spatial autocorrelation (and a near absence of negative spatial autocorrelation) exists in all maps. Further, the distribution of local Moran’s I demonstrates considerable heterogeneity in the spatial autocorrelation within each map. For example, in the T1w/T2w map, we see widespread elevated spatial dependencies in early somatosensory and visual regions as well as a range of anterior temporal and frontal regions. **Figure 2C** presents pairwise correlations between local Moran’s I across maps. Spearman’s rank correlation varies in amplitude and range from negative to positive for the different pairs of maps, reflective of varying alignment in spatial heterogeneity across different map pairs. In other words, in these maps of macroscale cortical properties, there is evidence that (i) within a map, spatial heterogeneity varies across the cortex; and the alignment of this heterogeneity varies across the brain maps.

### A 1-dimensional illustration of the problem

To understand the effects of these types of heterogeneity on spatial permutation approaches such as spatial randomization tests (Alexander-Bloch et al., 2018; Váša et al., 2018), we next consider some simulated datasets. Note that the objective of these simulations is to provide a simple illustration of the potential problem, rather than to model the type of stochasticity and signals in empirical brain maps. To this end, we first simulate a pair of 1-dimensional spatial signals with simple spatial autocorrelation which is heterogeneous over space. To make each signal, two different length 1D vectors drawn from a Gaussian distribution were generated and then resampled to create vectors with different amounts of spatial smoothness. We subsequently concatenate these vectors, creating a 1D vector that varies in spatial autocorrelation.

Figure 3. illustrates two scenarios with pairs of heterogeneous signals (represented using dashed and solid lines). These signals were both generated from random Gaussian noise independent from each other; therefore, a statistical test quantifying the similarity between each pair of signals should fail to reject the null hypothesis that the signals are independent. In one of the two scenarios **(Figure 3A)**, the heterogeneity is spatially aligned *i*.*e*., high or low spatial autocorrelation is present in the same locations in both signals. In the second scenario **(Figure 3B)**, the heterogeneity is inversely aligned. These scenarios are directly comparable to relationships between the autocorrelation of pairs of empirical maps, quantified by correlations between their local Moran’s I. The first simulated scenario corresponds to positive correlations between local Moran’s I, analogous to the comparison of the T1w/T2w ratio and cortical thickness. The second simulated scenario corresponds to negative correlations between local Moran’s I, analogous to the comparison of the T1w/T2w ratio with the evolutionary expansion map (see **Figure 2C**).

**Figure 3.**
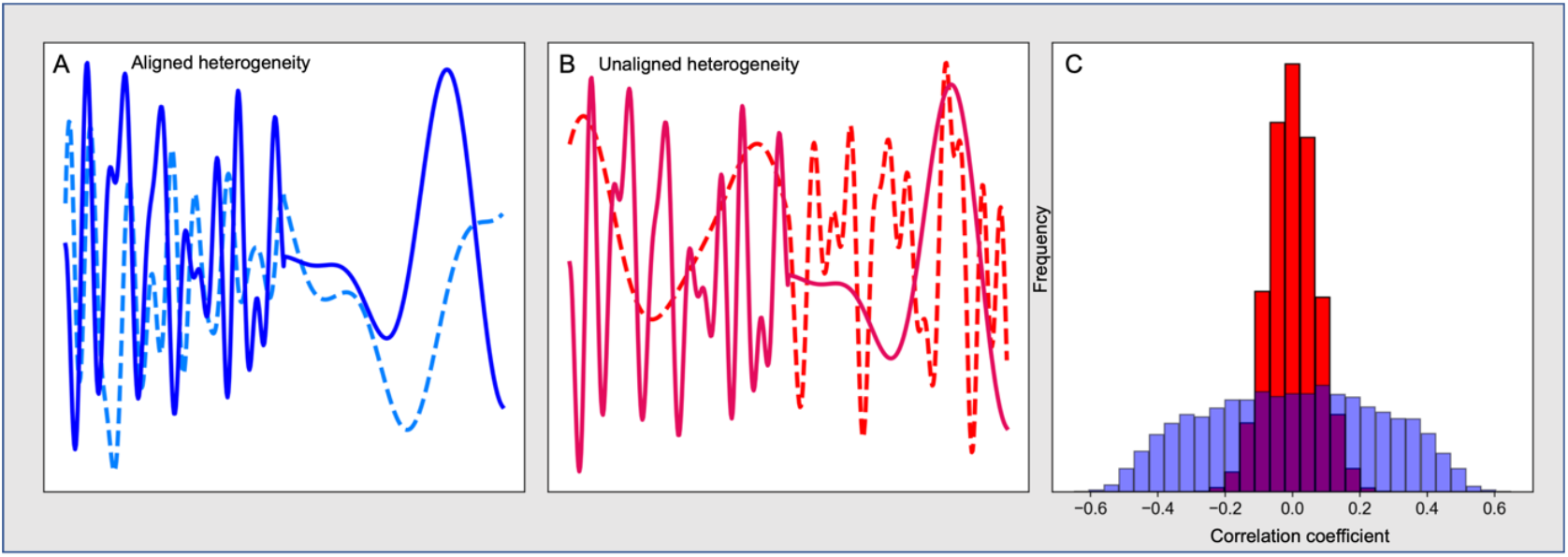
An illustration of different forms of heterogeneity in 1D spatial autocorrelation. Two random signals created by spatially concatenating vectors with high and low levels of smoothing are compared (dashed and solid lines). A) A case in which the heterogeneity of the two signals is aligned. B) A case where the heterogeneity of the two signals is inversely aligned. C) The distribution of pairwise Spearman’s rank correlation coefficients for the two scenarios; blue are aligned, red are unaligned (from 10,000 repetitions with different random seeds).

In **Figure 3C** we observe the distribution of Spearman’s Rank correlation coefficients between both variables, for 10,000 repetitions of the data generation process. Both for the aligned and non-aligned scenarios, we observe symmetric distributions of correlation coefficients centered around 0 (as expected because there is no correlation between the variables). However, the non-aligned scenario (high smoothness paired with low smoothness) shows a relatively narrow range of correlations, while the aligned variant shows a much wider distribution including more extreme correlation values. The difference between these two scenarios results from the spatial translation of one of the variables in relation to the other. Therefore, spatial translation applied to one of the signals (as is done with “spin” spatial randomization to build a null distribution) may result in null distributions with different statistical properties to the empirical result (for example, more extreme values). This can potentially lead to inappropriate statistical inference; whether it will inflate or deflate the probability of a false positive depends on the nature of the alignment of the heterogeneity across the true signals compared to the translated signals.

### A surface-based illustration of the problem

Above we have observed that the alignment of heterogeneity in the smoothness of a 1D signal can profoundly affect the distribution of correlations. To illustrate how this may affect statistical inference following spatial randomization tests on the sphere (commonly used for surface-based neuroimaging analyses; Alexander-Bloch et al., 2018; Váša et al., 2018), we created two additional simulated ground-truth datasets on a spherical surface. In one scenario, the data was generated by a spatially homogeneous process, where we anticipated that the spin permutation test would lead to appropriate statistical inference. In the other scenario, the data was generated by a spatially heterogeneous process, a 2D spherical analogue of the scenario in **Figure 3A**, which would be challenging for the spatial randomization test. For the heterogeneous situation, we used the method of Ingebrigtsen et al. (2014) to adjust the variance and length scale of the autocorrelation depending on the spatial location (**Figure 4A**). For each scenario, we created 5000 pairs of spheres using different random seeds. We then used the spin permutation procedure (with 100 iterations) (Alexander-Bloch et al., 2018; Markello and Misic, 2021) to generate a null distribution of correlations and obtain a measure of statistical significance as the proportion of null correlations which exceeds the true correlation coefficient. We repeated this process 5000 times for both the homogeneous and heterogeneous scenarios, resulting in a distribution of “true” correlation coefficients (between each pair of unpermuted maps) and a distribution of associated p-values derived from the spin permutations, for each scenario.

**Figure 4.**
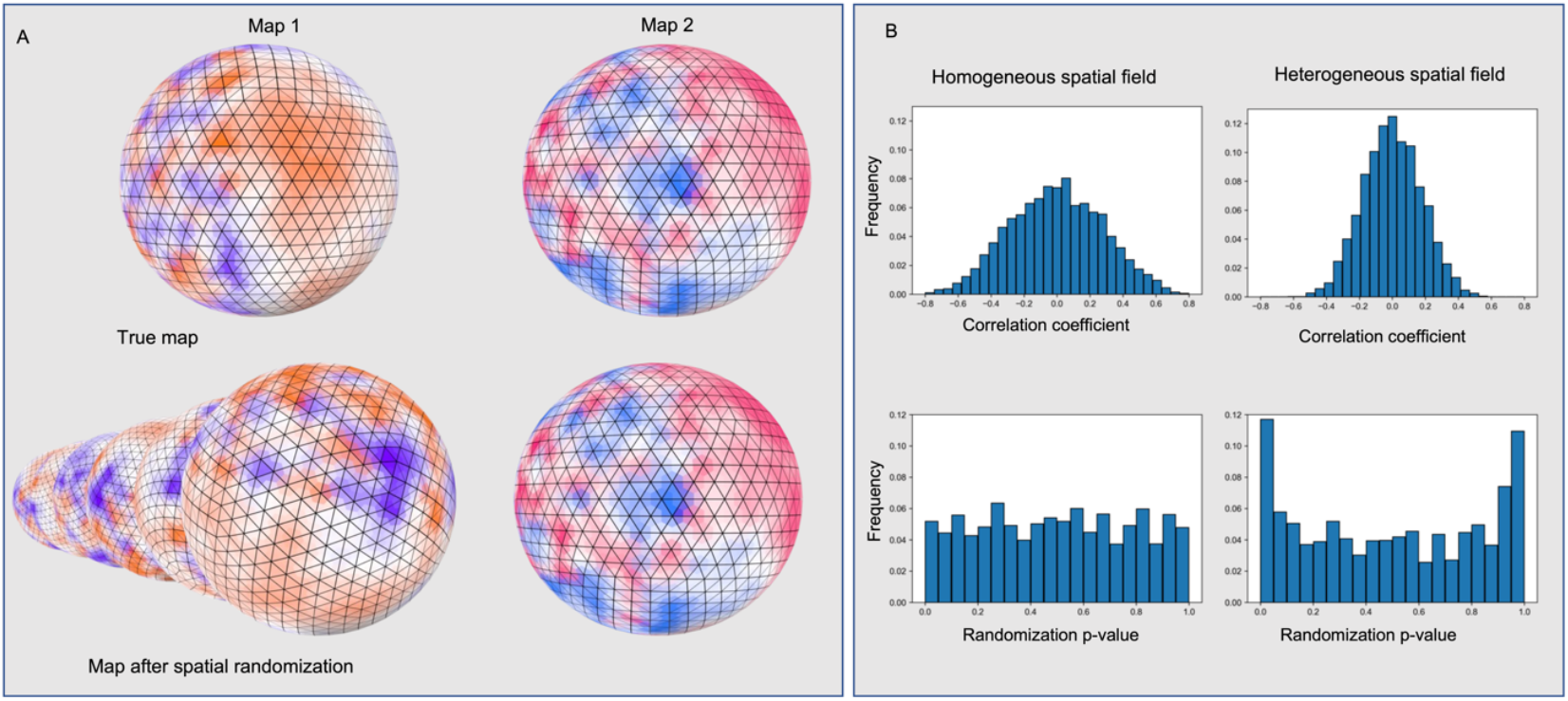
The impact of spatially heterogeneous autocorrelation on the spatial randomization test. **A**) Top: an illustration of two randomly generated maps with spatially-varying autocorrelation (created from a realisation of a stochastic partial differential equation on an icosphere surface). Bottom: one of these maps was randomly rotated 100 times to build a permutation distribution. In the heterogeneous scenario, the spatial autocorrelation varies based on the spatial location on the sphere (with different spatial autocorrelation regimes on either side of the 0 degree meridian). Pairs of these maps were generated with different random values and only aligned in terms of the heterogeneity of the spatial autocorrelation. Two scenarios were considered, homogeneous and heterogeneous spatial autocorrelation across the surface. **B**) Top: distribution of 5000 pairs of maps were generated and then spatially correlated (i.e., correlating the values across Maps 1 and 2), to generate “true” Spearman Rank correlation values. Bottom: Subsequently one map from each pair was randomly rotated 100 times to generate null distributions of correlations and associated p-values (defined as the proportion of null correlations exceeding the true correlation coefficient).

**Figure 4B** illustrates the distribution of correlation coefficients resulting from spatially correlating pairs of maps in both the homogeneous and heterogeneous scenarios, as well as the associated distribution of “spin” spatial randomization p-values. We observe that the homogeneous scenario results in a wider distribution of correlation coefficients; this is because it was generated with only a high level of spatial smoothing, while the heterogeneous scenario was generated with a mix of high and low spatial smoothing. For the homogeneous scenario, although the amount of spatial autocorrelation is higher resulting in more extreme correlation coefficients, the spatial randomization test works appropriately, controlling the error rate, with an approximately flat distribution of randomization p-values. In contrast, the heterogeneous scenario has a narrower range of correlation coefficients, but the spatial randomization test fails to adequately control the type 1 error, with a disproportionate number of extreme p-values compared to the null hypothesis. In this scenario, approximately 1/4 of p-values are within bins at the extremes of the histogram, compared to the expected 1/10 in the homogeneous situation. As such, any analysis involving comparisons between brain maps with heterogeneous spatial autocorrelation is likely to result in substantial inflation of p-values; the precise amount of inflation (or deflation) will depend on the degree of heterogeneity, the amount of spatial autocorrelation and the spatial resolution of the data.

#### Explicitly modelling spatial dependencies in the data

An alternative framework for statistically assessing the relationship between spatial maps in neuroimaging involves attempting to explicitly quantify the spatial autocorrelation while modelling the relationship between these maps. Several approaches have been developed in other scientific fields to this end, including R-INLA (Integrated Nested Laplace Approximation in R) (Lindgren et al., 2011; Lindgren and Rue, 2015; Bachl et al., 2019) and sdmTMB (Anderson et al. 2024). R-INLA uses approximate Bayesian methods to estimate a model of a continuous spatial process with stochastic partial differential equations realised on a discrete mesh. It allows for the creation of a generative model of spatial data, including various types of non-stationary spatial phenomena, as well as spatial covariates. Model comparison, using information criteria and/or out-of-sample cross-validation, enables differentiating between different hypotheses about the role of spatial covariates and assessing the spatial autocorrelation of the signal.

We first use R-INLA with a ground-truth model generated by a non-stationary process on a mesh (see **Figure 5**) generated with the same procedure as used in **Figure 4** above. Two maps (Map 1 and Map 2) were generated at 1000 random spatial locations on the sphere. Maps have different random seeds but the same non-stationary autocorrelation structure, with a binary variable based on the map’s *x*-coordinate included in the model’s dependency structure (see *Methods* and *Code* for full details). Three different models were then fit to the generated data, predicting Map 1 with: *(i)* a stationary Gaussian random field as the intercept; *(ii)* a stationary field plus Map 2 as a covariate; *(iii)* a non-stationary Gaussian field as intercept (similar to the one used to generate the data). Model comparison was performed using the Deviance Information Criterion (Spiegelhalter et al., 2002) as well as root-mean-squared-error (RMSE) comparing predicted and actual values of Map 1 at 5000 out-of-sample coordinates on the sphere. For both DIC and RMSE, R-INLA correctly identifies the non-stationary model without a covariate for Map 2, as the best model (which is consistent with how the data was generated). Note that the models with stationary fields (with and without a covariate) are much more similar (in terms of both RMSE and DIC).

**Figure 5.**
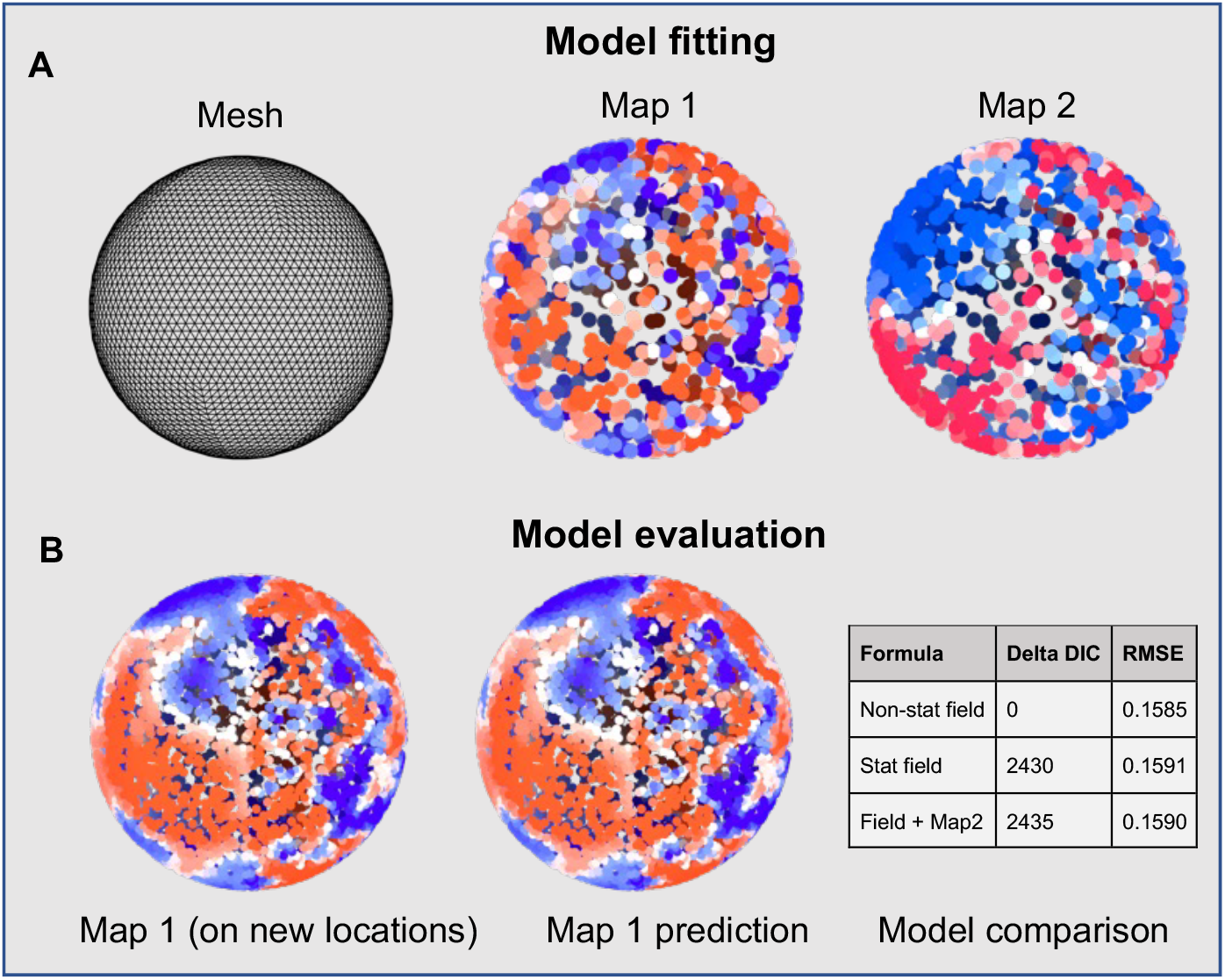
Explicitly modelling spatial autocorrelation in a simulated dataset on a sphere, generated by a non-stationary process. **A)** Model fitting: a spherical mesh is approximated, and a pair of maps generated at random locations on the sphere; then, the mesh is used to fit three different models to predict Map 1: (i) a stationary spatial field; (ii) a stationary spatial field plus Map 2; (iii) a non-stationary spatial field. **B)** The best model is selected by evaluating model performance on either the deviance information criteria (DIC) or the root mean squared error between predicted and actual values of Map 1 at out-of-sample spatial locations.

The above simulated example illustrates an alternative approach for studying the relationship between different cortical brain maps. We illustrate this by comparing the spatial relationship between the principal functional gradient (Margulies et al. 2016) and a T1w/T2w based proxy for intracortical myelin (Glasser et al. 2011) (**Figure 6A**). We also illustrate how explicit modelling of the spatial process using a range of different models can account for different types of spatial relationships between the maps (see **Figure 1**). In particular, we can evaluate different non-stationary contributions to spatial autocorrelation (i.e., spatial variation in the correlational structure between the maps, not just the spatial dependency structure within a map), including maps with spatially-varying coefficients, where instead of a single coefficient being used to predict the whole map (e.g., in spatial ordinary least squares regression), the coefficient can vary across the cortex.

**Figure 6.**
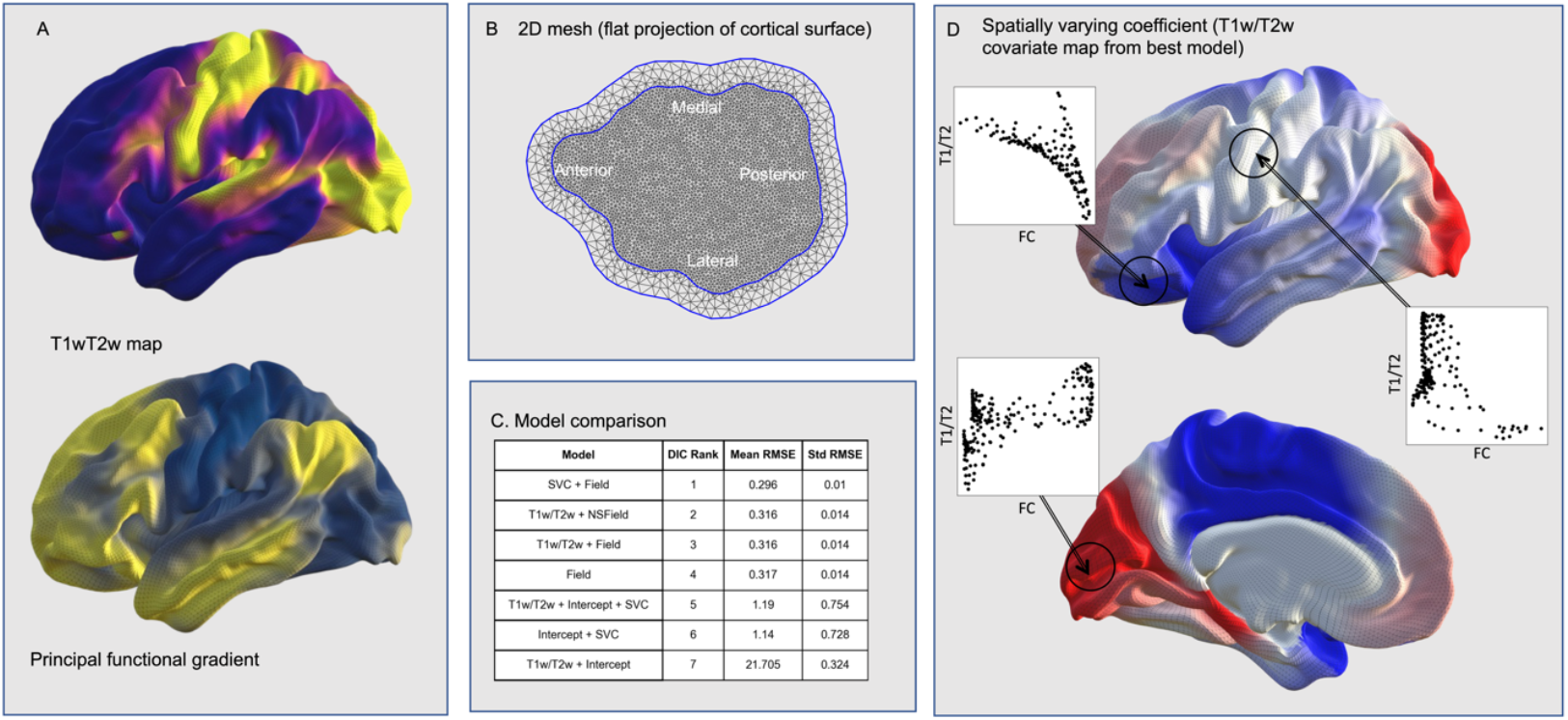
Comparing two empirical cortical maps by explicitly modelling spatial processes. **A)** The maps being compared: the T1w/T2w ratio (an approximate measure of intracortical myelin) and the principal functional gradient. **B)** A mesh made from a subset of the data projected onto a plane, including an outer layer to control for potential boundary effects. This mesh was used to fit models on a different subset of the data from both T1w/T2w and the functional gradient. **C)** Model comparison reveals that a spatially varying coefficient (T1w/T2w) model with a stationary field has the lowest Deviance Information Criterion (DIC) and mean out of sample error (RMSE) across 10-fold cross-validation. **D)** The spatially varying coefficient model reprojected back onto the cortical surface (cold/warm colors indicate a negative/positive relationship between T1w/T2w and the principal functional gradient). Insets display scatterplots of the principal functional gradient values with the T1w/T2w values within illustrative regions of interest.

We first split the data into 10 folds, with 90% of the data available for model fitting, and 10% held out for subsequent assessment of model fit for each fold. For each fold, a random proportion of the data (10% of all vertices excluding the medial wall: approximately 3000 vertices) is used to make a mesh using the *fmesher* tool (Bachl et al., 2019). For simplicity, we use a flat projection of the cortex, although it is also possible to use other projections to make the mesh. Subsequently, a range of different models were fit to a random 30% of the remaining data. We used a restricted set of the data for computational efficiency, although the analysis is also feasible on the full dataset (see Github repository). We used a non-exhaustive selection of models accounting for multiple spatial dependency scenarios. In our simulation, the principal gradient map (Margulies et al. 2016) was predicted by each of the following models:

(i) A linear relationship with T1w/T2w and constant intercept with no Gaussian random fields (T1w/T2w + intercept);
(ii) A Gaussian random field (instead of a constant intercept) and no covariate (Field);
(iii) A Gaussian random field plus the T1w/T2w map (T1w/T2w + Field);
(iv) A non-stationary field (instead of a constant intercept) with the T1w/T2w map as the explanatory covariate in the dependency structure and an additional constant T1w/T2w covariate (T1w/T2w + NSField);
(v) A spatially varying coefficient (T1w/T2w) plus a Gaussian random field (instead of a constant intercept; SVC + Field);
(vi) A spatially varying coefficient (T1w/T2w) with a constant intercept (SVC + Intercept);
(vii) A spatially varying coefficient (T1w/T2w) with a constant intercept and a constant T1w/T2w covariate (T1w/T2w + Intercept + SVC).

Model selection was performed both with Deviance information criteria on the fitted model and out-of-sample prediction on 10% of the data that was held out from mesh creation and model fitting for each fold. (Note that the whole dataset was not used in the interest of computational efficiency.)

Based on both the Deviance Information Criteria and out-of-sample RMSE, the spatially-varying-coefficient and Gaussian field model performed best for every fold (See **Figure 6C**). This suggests that there is a relationship between the principal functional gradient derived from rs-fMRI and the T1w/T2w map, but that it is spatially varying. The spatially-varying coefficients can be projected onto the cortical surface (**Figure 6D**). A range of qualitatively different relationships can be seen across cortex, ranging from predominantly negative relationships centered around the insula and paracentral lobule, through a lack of relationship in in a subset of regions including primary motor cortex, to positive relationships centered around the occipital pole (see **Figure 6D;** this can also be observed with scatter plots of in the raw data in specific regions, see insets in **Figure 6D**).

## Discussion

Due to the increasing recognition of the importance of macroscale properties of the brain as a determining factor of its function (Margulies et al., 2016, Smallwood et al., 2022), many studies have assessed statistical similarities between brain maps (e.g., Larivière et al., 2021, 2023; Markello et al., 2022; DeKraker et al., 2024). Here, we illustrate that these statistical comparisons depend upon correctly modelling the heterogeneity in the spatial autocorrelation of the signal *within* individual maps. In other scientific fields, a range of techniques have been developed for modelling spatial processes to characterize spatial maps and their relationships. Here, we illustrate that one such approach, based on modelling spatial processes as stochastic partial differential equations (Lindgren et al., 2011), explicitly models spatial heterogeneity and so provides a framework for correctly evaluating similarities between different brain maps.

As well as supporting more accurate statistical inferences, explicitly modelling spatial processes also provides the opportunity for more sophisticated descriptions of spatial phenomena. Here, we saw that over and above assessing for the presence of a spatial relationship between maps, we can also observe whether that relationship is spatially varying; that is, quantify whether the sign and amplitude of the relationship vary according to a spatial process, rather than a single spatial correlation value across the brain. This may capture more biologically realistic relationships than a single correlation value; for example, we can allow the effects of a neurotransmitter receptor map (e.g., 5-HT2A) to relate differently to functional activity in different areas of the cortex or subcortical regions, in a single statistical model that also accounts for spatial autocorrelation.

Another advantage of this approach is that it can be used to incorporate multiple replicates within a single spatial model. For example, non-stationarity in a spatial process can reflect variation in the mean or the covariance structure within a map; this difference can only be differentiated with multiple replicates (e.g., multiple different maps within or across individuals). It can also accommodate group differences, e.g., investigating how the relationship between spatial maps changes with patient and control groups, and allows for quantifying changes in the spatial processes (e.g., the covariance structure) across groups. We can also extend the model to consider multiple covariates simultaneously (e.g., Tavor et al., 2016), for example to assess how a range of neurotransmitter maps or microstructural features reflect functional properties such as the principal functional gradients, with different random Gaussian fields for the different covariates, (including potential interactions between covariates). Such statistical approaches can help model and disentangle the likely complex spatial relationships between these maps, and between the underlying genetic, developmental, anatomical and physiological processes. In a related vein, spatio-temporal models can also be realised, modelling a temporal autoregressive process in addition to the spatial one, allowing for a more comprehensive perspective on neural dynamics (Mejia et al., 2020), including task-evoked signal change. A similar spatio-temporal approach can also be applied to study resting state functional data without having to first apply techniques such as pairwise correlation or independent component analysis to integrate across the temporal dimension).

While approaches to explicitly model the underlying spatial processes can better characterize statistical relationships between spatial maps, the added flexibility carries a cost. Whereas approaches such as spatial randomization tests (Alexander-Bloch et al., 2018; Váša et al., 2018) are easy to apply, using R-INLA or other approaches to model spatial processes within brain maps requires domain knowledge to inform the selection of model priors and choice of covariates. Also, results may be more ambiguous and harder to interpret than the resulting *r* and *p*-values from null-modeling approaches. It is also worth emphasizing that the existing null model approaches are still likely to be appropriate in a wide range of scenarios, for example, if the compared maps exhibit relatively limited spatial autocorrelation, and this is relatively spatially homogeneous. Finally, approaches such as R-INLA have been developed primarily for 2D or spherical maps; in future, it would be beneficial to build 3D meshes to more accurately model the cortical surface, and to consider approaches which can be applied to 3D volumes.

In summary, variability in the spatial autocorrelation within cortical maps poses challenges for statistical comparisons between them using existing null model frameworks. Alternative methods such as stochastic partial differential equations offer a more comprehensive approach to modeling and comparing spatial maps, accommodating different forms of spatial heterogeneity.

## Methods

### Neuroimaging data

All neuroimaging data were obtained from *neuromaps* (Markello et al., 2022) in the FSLR surface projection. We used four different maps from three different studies. The T1w/T2w maps and the cortical thickness maps are the group averages from the Human Connectome Project’s Young Adult 1000 participant release (Glasser et al., 2013). The principal gradient of functional connectivity was calculated on the resting state fMRI data from the HCP Young Adult dataset using diffusion embedding (Margulies et al., 2016). The evolutionary expansion map was created by Hill et al. (2010). For full details of the pre-processing pipelines and subsequent creation of each map, see the corresponding publications.

### Local Moran’s I

Code to calculate local Moran’s I was adapted from Markello and Mišić (2021). For computational reasons, the four cortical maps were downsampled to 10,000 vertices; the distance matrix was calculated based on the down-sampled version of the FSLR atlas (with 5000 vertices in each hemisphere). Local Moran’s I was calculated for each vertex in each of the four maps.

### One-dimensional simulation

Code to reproduce the 1D scenario is available on Github (link below). Two independently and identically distributed arrays were created with: (i) 5 samples; (ii) 100 samples. Both were then interpolated with the *scipy* resample function to 1000 samples. These two 1000-sample arrays were normalized (demeaned and then divided by the standard deviation) and then concatenated to form a 2000-sample 1D array with both a high and low smoothness regime. Pairs of 1D arrays were created with different random seeds. Differences in the order of the concatenation (high-low or low-high) across each of the pairs determined whether the heterogeneity of smoothness was aligned or not. Finally, the pairs of arrays were correlated. This was repeated 10,000 times to build distributions of correlation coefficients for aligned and inversely aligned smoothness.

### Surface-based simulation on a sphere

Spherical spatially heterogeneous and homogeneous spatial autocorrelation was generated using R-INLA (Lindgren and Rue, 2015) and the INLABRU (Bachl et al., 2019) wrapper package. The full code is available on GitHub. A spherical mesh was generated from an icosphere with 30 subdivisions (using the *fmesher* library). A binary variable was created for each point on the mesh, based on its *x*-coordinate; this was then demeaned. A model of a Gaussian random field with a Matérn covariance kernel was instantiated on the mesh; either a standard homogeneous Gaussian random field or a spatially varying Gaussian random field (based on the binary variable) were used. Five thousand pairs of sample maps were created with different random seeds. Full details of the model parameterisation are in the associated R code (Krainski et al., 2018). To assess spin test performance, these pairs of random maps with either homogeneous or aligned spatially varying autocorrelation structure were compared using code from Markello and Mišić (2021).

In addition, a separate simulation was performed in Python (see GitHub) by explicitly smoothing points on the sphere based on two smoothing kernels, cutting them in half, normalizing them, and then adding them together to create aligned heterogeneity (similar to the 1D simulated data above). This led to qualitatively similar results to the simulated data generated from the R-INLA model.

### R-INLA modelling of spatial processes

#### (i) Simulated data

A fine spherical mesh (icosphere with 30 subdivisions) was created with the same model used to generate heterogeneous spatial maps for evaluating the spin test. Two randomly-seeded maps were then generated (with an identical spatial process) at 1000 random locations on the sphere (not on the mesh). A new lower-resolution mesh was created (again an icosphere, but with 20 subdivisions). Three models were fit to the data, predicting Map 1, with either: *(i)* A stationary random Gaussian field; *(ii)* A stationary random field and Map 2 as a covariate; *(iii)* Or a spatially varying random Gaussian field (with the hemisphere controlling the spatial variability in the field). Model performance was assessed with two metrics: (a) the Deviance Information Criterion (Spiegelhalter et al., 2002) on the trained data; and (b) out of sample root-mean-squared error on 5000 novel randomly selected locations on the sphere.

#### (ii) Cortical maps

The group average principal functional gradient (Margulies et al., 2016) and the T1w/T2w map (Glasser et al. 2013) were used. The 2D projection was the default flat map from the HCP Young Adult group average data release. The model was fit with 10-fold cross validation: different subsets of the training data (i.e., different spatial locations) were used for mesh creation, and a separate set was used for model fitting for each fold. Model evaluation again used both the Deviance Information Criteria and root mean squared error on held out 10% of the data used for evaluation at each fold. Within each fold the 2D mesh was generated from 10% of the data using the *fmesher* library. The same 7 models (listed in the results) were fit to each fold assessing combinations of linear effects, stationary and non-stationary Gaussian random fields, and spatially varying coefficients. For non-stationary Gaussian random fields, the demeaned T1w/T2w ratio was used as a covariate in the formula for the field. We used a restricted set of the data for computational efficiency reasons and to minimize data leakage (since the analysis was repeated across multiple folds). One example on the GitHub repository fits the INLA model to a much larger portion of the data, illustrating that it is computationally feasible.

Code to repeat the analyses is openly available at: https://github.com/ActiveNeuroImaging/SpatialNonStationarity

